# Availability of charged tRNAs drives maximal protein synthesis at intermediate levels of codon usage bias

**DOI:** 10.1101/2025.06.16.659965

**Authors:** Alexis M. Hill, Kelly To, Claus O. Wilke

## Abstract

Synonymous codon usage can influence protein expression, since codons with high numbers of corresponding tRNAs are naturally translated more rapidly than codons with fewer corresponding tRNAs. Although translation efficiency ultimately depends on the concentration of aminoacylated (charged) tRNAs, many theoretical models of translation have ignored tRNA dynamics and treated charged tRNAs as fixed resources. This simplification potentially limits these models from making accurate predictions in situations where charged tRNAs become limiting. Here, we derive a mathematical model of translation with explicit tRNA dynamics and tRNA re-charging, based on a stochastic simulation of this system that was previously applied to investigate codon usage in the context of gene overexpression. We use the mathematical model to systematically explore the relationship between codon usage and the protein expression rate, and find that in the regime where tRNA charging is a limiting reaction, it is always optimal to match codon frequencies to the tRNA pool. Conversely, when tRNA charging is not limiting, using 100% of the preferred codon is optimal for protein production. We also use the tRNA dynamics model to augment a wholecell simulation of bacteriophage T7. Using this model, we demonstrate that the high expression rate of the T7 major capsid gene causes rare charged tRNAs to become entirely depleted, which explains the sensitivity of the major capsid gene to codon deoptimization.

## 1 Introduction

Many fast growing, unicellular organisms maintain imbalanced tRNA copy numbers, such that for each isoaccepting group of tRNAs, one tRNA species is often present in higher cellular concentrations than the other(s) (Ikemura 1981a,b, 1982). From an evolutionary perspective, the cause of imbalanced tRNA copy numbers remains an open question (Plotkin and Kudla 2011). However, one consequence is that individual codons can be translated at different rates depending on whether their cognate tRNAs are more or less common in the cellular tRNA pool (Sørensen et al. 1989). By changing synonymous codon frequencies so that they are more optimal with respect to host-cell tRNA abundances, it is possible to improve heterologous protein expression rates (Gustafsson et al. 2004; Welch et al. 2009). Codon modification has also been used as a strategy to attenuate viruses (for vaccine development) (Bull et al. 2012; Burns et al. 2006; Coleman et al. 2008) and to improve protein co-translational folding (Liu 2020; Walsh et al. 2020), and in general, codon usage is considered to be an important determinant of translation efficiency. Computational models that make precise predictions about the relationship between codon usage and gene expression could potentially improve our ability to rationally design re-coded organisms with desired biological properties.

The relationship between codon usage bias and protein production has been modeled extensively, using different mathematical approaches and at different levels of granularity (Choi and Covert 2023; Cope and Shah 2025; Dykeman 2024; Katz et al. 2022; Levin and Tuller 2020; Raveh et al. 2016; Reuveni et al. 2011; Seeger et al. 2023; Shah and Gilchrist 2011; Shah et al. 2013; Zarai et al. 2016). Many existing models do not explicitly model tRNA aminoacylation (charging) even though translation speeds ultimately depend on the concentration of charged tRNAs (Sørensen et al. 1989). Under steady-state conditions, assuming constant charged tRNA availabilities is likely a reasonable simplification since most types of tRNAs are are around 80% charged at any given time (Dittmar et al. 2005), and the overall composition of the tRNA pool does not meaningfully change on short time scales (Dittmar et al. 2004; Jakubowski and Goldman 1984). However, under conditions of environmental stress (such as amino acid starvation) pools of charged tRNAs can rapidly rearrange or even be entirely depleted (Dittmar et al. 2005; Dong et al. 1996; Sørensen 2001; Torrent et al. 2018). Models that treat charged tRNAs as fixed resources may fail to accurately capture non-equilibrium gene expression dynamics.

In a recent study, we introduced a stochastic model of translation that incorporates explicit tRNA dynamics (Roots et al. 2025), where charged tRNA availability is dependent on the interplay between ribosome activity, codon usage, and tRNA re-charging. The model was used in combination with fluorescent protein expression experiments to argue that codon deoptimization (the replacement of preferred codons with non-preferred codons) reduces protein expression, that the extent of this effect is modulated by the degree to which codon usage bias matches the available tRNA pool, and that there can be a regime of overoptimization where the trend reverses and where codon optimization (the replacement of non-preferred codons with preferred codons) similarly leads to reduced protein expression. In aggregate, Roots et al. (2025) suggested that dynamic tRNA pools are critical for an accurate description of the effect of codon usage bias on protein expression. Here, we take this work further by developing a mathematical model of this system that can be solved numerically, without the need for stochastic simulations, allowing for much more rapid and systematic exploration of the model’s parameter space. We also apply the insights gained from this model to the case of viral attenuation via codon deoptimization in T7. Our modeling results suggest that non-preferred charged tRNAs get fully depleted during the final stages of the T7 replication cycle, even for wild-type T7. This causes a substantial fitness penalty when the most highly expressed T7 gene gets recoded to use a much higher fraction codons corresponding to these depleted tRNAs.

## 2 Results

### 2.1 Mathematical model

Roots et al. (2025) used a simplified model of codon usage where there are two synonymous codons and two corresponding species of tRNAs (Fig. 1). The two species of tRNAs in the model are allowed to have different abundances, and the more abundant tRNA is referred to as the *preferred tRNA*. Similarly, the codon that corresponds to the preferred tRNA is referred to as the *optimal codon*, and the other codon as the *non-optimal codon*. The total numbers of tRNAs and ribosomes are held constant over time, so that only the fractions of charged/uncharged tRNAs and bound/unbound ribosomes change dynamically in this system. Finally, the transcript population in this model is homogeneous, such that the strength of ribosome binding, the codon usage frequency, and the length of each transcript are the same across all transcripts.

**Fig. 1.**
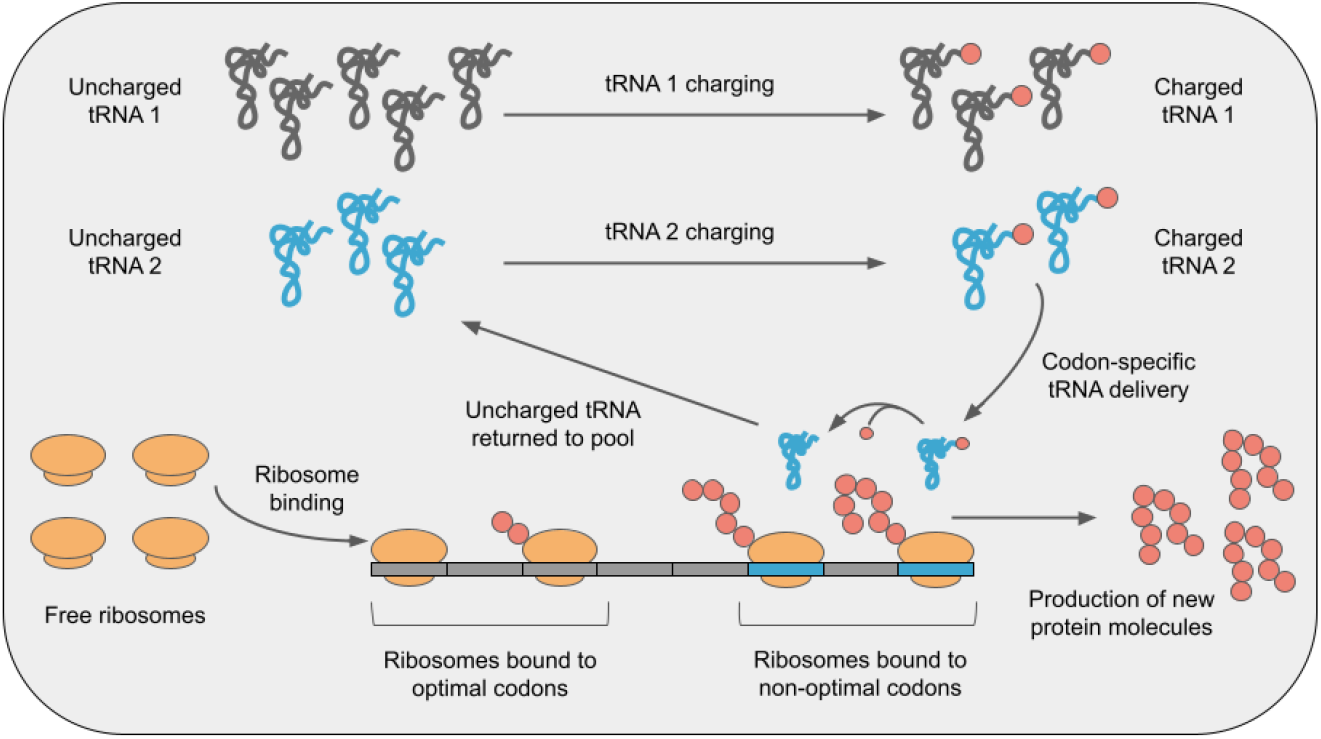
Schematic of the two-species tRNA model considered here. tRNAs can be either uncharged or charged, and each tRNA species is recharged by its own charging reaction. Ribosome elongation from any individual codon requires at least one charged tRNA corresponding to the codon the ribosome is bound to. Elongation converts charged tRNAs into uncharged tRNAs, which are unable to participate in translation until being recharged.

We will derive a set of differential equations that describe the dynamics of this system. First, for every tRNA species in this system, uncharged molecules are converted into charged molecules with rates *k*_charge_*T*_u*i*_, where *k*_charge_ is the tRNA charging rate and *T*_u*i*_ is the number of uncharged tRNA molecules of type *i*. For simplicity, we assume equal *k*_charge_ for all tRNA charging reactions. The rate at which charged tRNA molecules are depleted by ribosome elongation from cognate codons is *k*_speed_*R*_b*i*_*T*_c*i*_, where *k*_speed_ is the ribosome elongation rate constant, *R*_b*i*_ is the number of ribosomes bound to codons of type *i* (in other words, the codon occupancy), and *T*_c*i*_ is the number of charged tRNA molecules of type *i*.

Let *T*_c1_ and *T*_c2_ be the abundances of charged preferred and non-preferred tRNAs, respectively. These abundances change over time according to

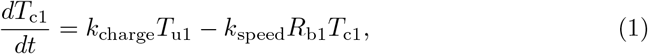

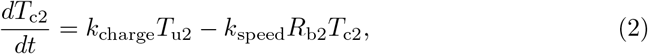

where *T*_u1_ and *T*_u2_ are the abundances of uncharged preferred and non-preferred tRNAs and *R*_b1_ and *R*_b2_ are the abundances of ribosomes bound to optimal and non-optimal codons, respectively.

Ribosomes bind to mRNAs dynamically with rate *k*_bind_*NR*_f_, where *k*_bind_ is the ribosome binding rate constant, *N* is the number of ribosome binding sites, and *R*_f_ is the number of free ribosomes. Note that because each transcript has one ribosome binding site, *N* also corresponds to the number of transcripts in the system. Ribosomes become unbound automatically after reaching the end of a transcript, which makes the rate of translation termination (the off-rate) proportional to the sum of each per-codon elongation rate and inversely proportional to the transcript length, *L* (in codons). Overall, free ribosome dynamics are governed by the trade-off between gain due to elongation/translation termination and depletion due to translation initiation. We can express this relationship as

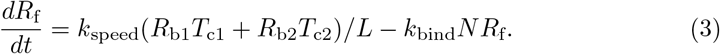

Finally, as ribosomes move along transcripts, they alternate between being bound to optimal and non-optimal codons. We assume that we can describe this movement via a mean-field approximation that considers only the relative fraction of codons of the two types but not their spatial arrangement along the transcripts. Let *f*_op_ be the fraction of optimal codons in a transcript. The rate at which ribosomes not already bound to optimal codons move and land on optimal codons (the conversion rate) is then given by *k*_speed_*R*_b2_*T*_c2_*f*_op_, that is, the rate of ribosome elongation from non-optimal codons times the fraction of optimal codons, *f*_op_. Similarly, for non-optimal codons, the conversion rate is *k*_speed_*R*_b1_*T*_c1_(1 *− f*_op_). The overall dynamics for bound ribosomes are then described by

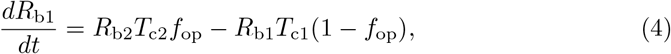

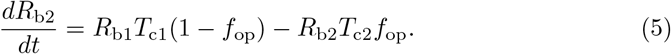

In steady state, all tRNA and ribosome abundances are constant. Therefore, we can calculate steady-state abundances by setting Eqs. (1)–(4) to zero. This results in the following expressions:

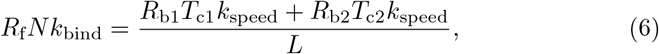

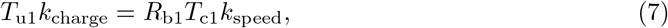

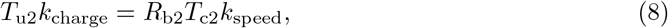

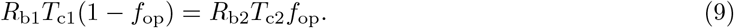

This system of equations is underdetermined, but we can arrive at a unique solution by considering that the species totals for ribosomes, *R*^tot^, and for the two types of tRNAs,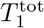, and 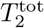, are constant and given by:

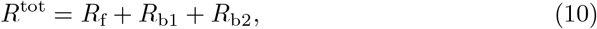

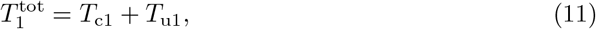

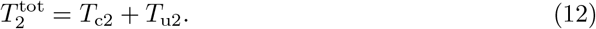

Eqs. (6)–(12) can be solved numerically to describe the steady-state behavior of our two-codon system with dynamic tRNA abundances.

We can calculate the protein expression rate *P*_r_ by considering that new protein molecules are produced as ribosomes reach the end of a transcript. Therefore, the protein expression rate is equal to the ribosome off-rate:

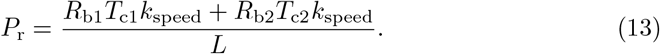

Here, we will take advantage of the substantially reduced computational cost of solving Eq. (13) versus running stochastic simulations to explore the relationship between tRNA dynamics, codon usage, and protein expression more systematically.

### Analysis of the two-codon system

We first verified that numerically solving Eq. (13) yields identical results to the previously published (Roots et al. 2025) stochastic simulations of the two-codon system.

In the stochastic simulations, an ensemble of identical transcripts was used to represent the background endogenous mRNA population; then, another smaller number of transcripts was added to represent an overexpressed exogenous gene. To simplify our analysis, here we removed the exogenous transcript species from the model, leaving only the transcripts representing an endogenous mRNA. We also reduced the transcript length of the endogenous mRNA from 1000 to 300 codons. We kept all other species counts and initial conditions as described (Roots et al. 2025). We found that the numerical solutions showed very good agreement with the simulation when solved using an identical set of parameters (Fig. S1).

With the mathematical model verified, we next used it to systematically explore the relationship between tRNA dynamics, codon usage, and protein expression. One free parameter in this model is the ratio of preferred to non-preferred tRNAs. In the prior work, a tRNA ratio of 7:3 (1750 preferred tRNAs and 750 non-preferred tRNAs) was chosen based on a back-of-the-envelope calculation of the average ratio of common and rare tRNAs across all isoaccepting groups of tRNAs, using measurements from *E. coli* (Dong et al. 1996). Here, we considered two additional tRNA ratios (one with more skew and one where tRNA abundances are equal) to explore how variation in tRNA abundances impacts protein expression. Using the model with equal tRNA abundances as the baseline, we calibrated the three remaining free parameters in the model (the rate constants for ribosome binding, *k*_bind_, ribosome elongation, *k*_speed_, and tRNA charging, *k*_charge_) to bacterial gene expression properties measured empirically (Table 1). We then systematically varied the tRNA charging rate constant to make tRNA re-charging more or less limiting relative to the baseline elongation rate for each of the three tRNA ratios. For each model parameterization, we solved Eqs. (6)–(12) numerically for the steady-state optimal and non-optimal codon occupancies (*R*_b1_ and *R*_b2_) and steady-state charged tRNA availability (*T*_c1_ and *T*_c2_), and then used Eq. (13) to calculate the protein expression rate, in molecules per second.

**Table 1.**
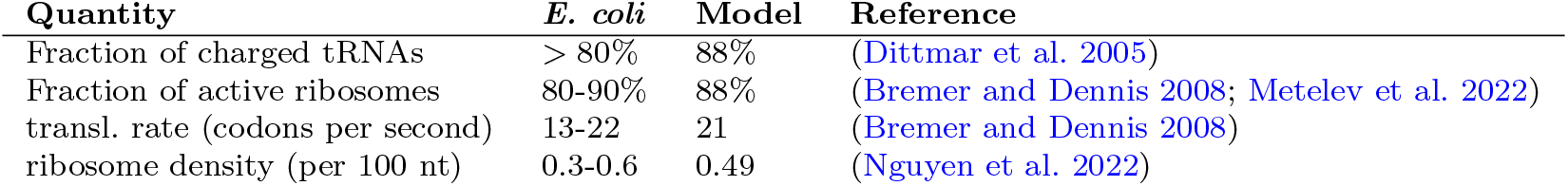
Steady-state translation properties in *E. coli* and in the two-codon model.

Our analysis revealed that the protein expression rate generally depends on the amount of codon usage bias, but the dynamic range in protein expression rates is influenced by both the amount of skew between the two tRNA species and the tRNA re-charging rate. For example, when there is more skew in tRNA abundances (fraction of preferred tRNAs = 0.9) and the tRNA charging rate is high, increasing the fraction of optimal codons from 0 to 1 increases the overall protein expression rate by at least 5-fold (Fig. 2A right-most panel). When the tRNA skew is more modest (fraction of preferred tRNAs = 0.7, Fig. 2A center panel) increases in gene expression rates are also more modest, and when both tRNAs are equally abundant (fraction of preferred tRNAs = 0.5, Fig. 2A left-most panel), protein expression is completely insensitive to codon usage at high values of *k*_charge_.

**Fig. 2.**
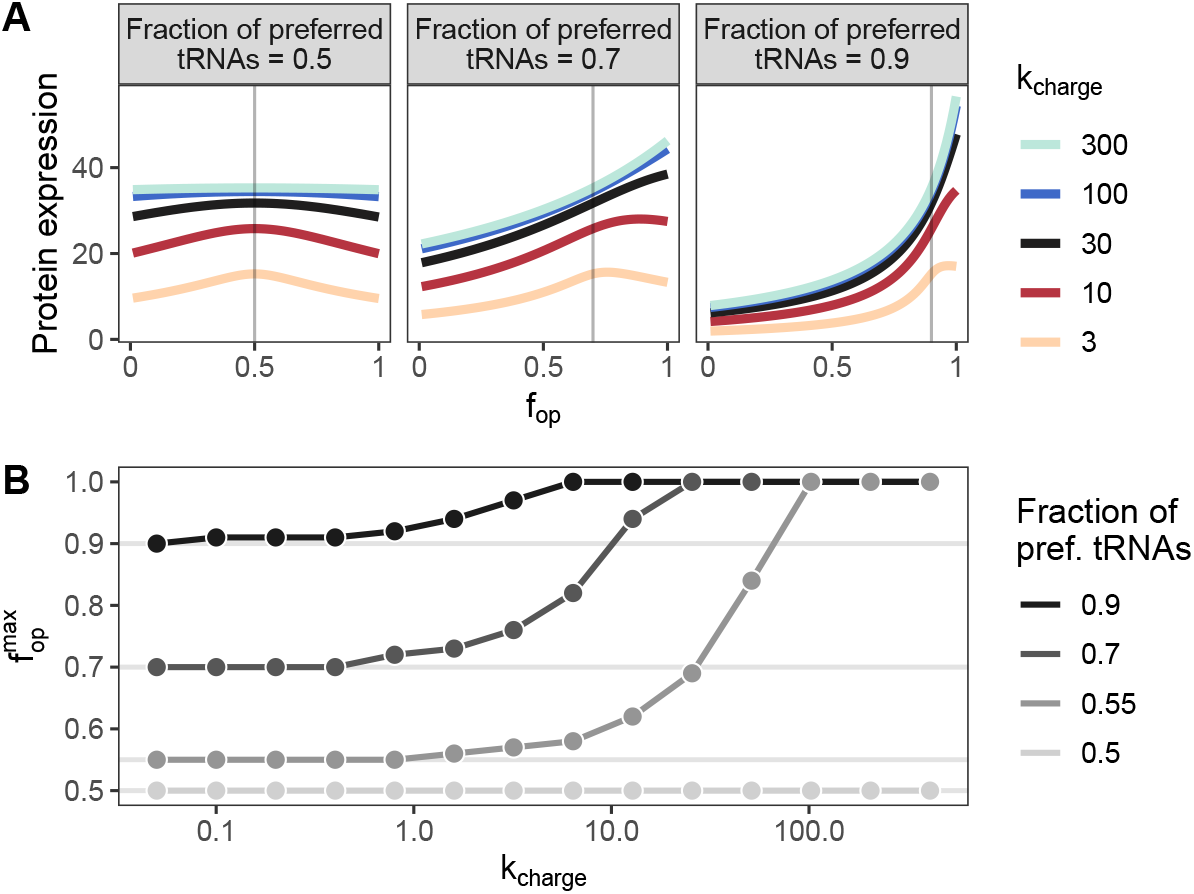
The effects of codon usage on protein expression. A: Protein expression rate, in molecules per second, as a function of the fraction of optimal codons, *f*op. Here, an *f*op of 0 corresponds to 0% optimal codon usage, and an *f*op of 1 corresponds to 100% optimal codon usage. The midpoint on the color scale (black) indicates simulations with the calibrated *k*_charge_. The gray vertical lines indicate the *f*op where the fraction of optimal codons matches the fraction of preferred tRNAs. B: Relationship between the *f*_op_ that maximizes protein expression, 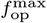, and *k*_charge_. For each group of simulations, as *k*_charge_ increases, 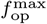 increases from a value that is exactly equal to the tRNA ratio (indicated by the gray horizontal lines) to 1.

A striking feature of the dependency of protein expression on *f*_op_ is the appearance of a maximum that seems, for small tRNA charging rates, to coincide with the fraction of preferred tRNAs, but tends to move towards larger *f*_op_ for higher charging rates (Fig. 2A). We thus systematically explored how this maximum depends on the fraction of preferred tRNAs and on *k*_charge_. We found that indeed the location of the maximum converged to the fraction of preferred tRNAs for low *k*_charge_ and to 1 for high *k*_charge_ (Fig. 2B). And, the closer the fraction of preferred tRNAs was to 0.5, the higher *k*_charge_ had to be for the location of the optimum to move. Finally, at a fraction of preferred tRNAs of exactly 0.5 the location of the optimum did not move at all (Fig. 2B). This latter observation is expected due to symmetry, as at a fraction of preferred tRNAs of exactly 0.5 the concept of preferred and non-preferred tRNAs is no longer meaningful. In this limit, both tRNA species behave exactly the same.

The finding that 100% optimal codon usage maximizes protein expression only when tRNA charging is high makes intuitive sense when we consider how the abundances of charged tRNAs respond to changes in codon usage under different *k*_charge_ regimes. At high rates of tRNA charging, both preferred and non-preferred tRNAs remain fully charged, regardless of codon usage (Fig. 3, blue curves). In practice, this means that uncharged tRNAs get replenished nearly instantaneously, i.e., before another ribosome elongation event occurs. In this case, using 100% optimal codons maximizes protein expression, because the corresponding charged preferred tRNA is always more readily available than the charged non-preferred tRNA. At the other extreme, for low tRNA charging rates, overuse of either codon depletes the charged tRNA for that codon, while, importantly, the other tRNA remains charged in excess of what is needed for translation (Fig. 3, orange curves). In this regime, balancing codon usage to the tRNA pool minimizes the number of excess charged tRNAs and maximizes the protein expression rate.

**Fig. 3.**
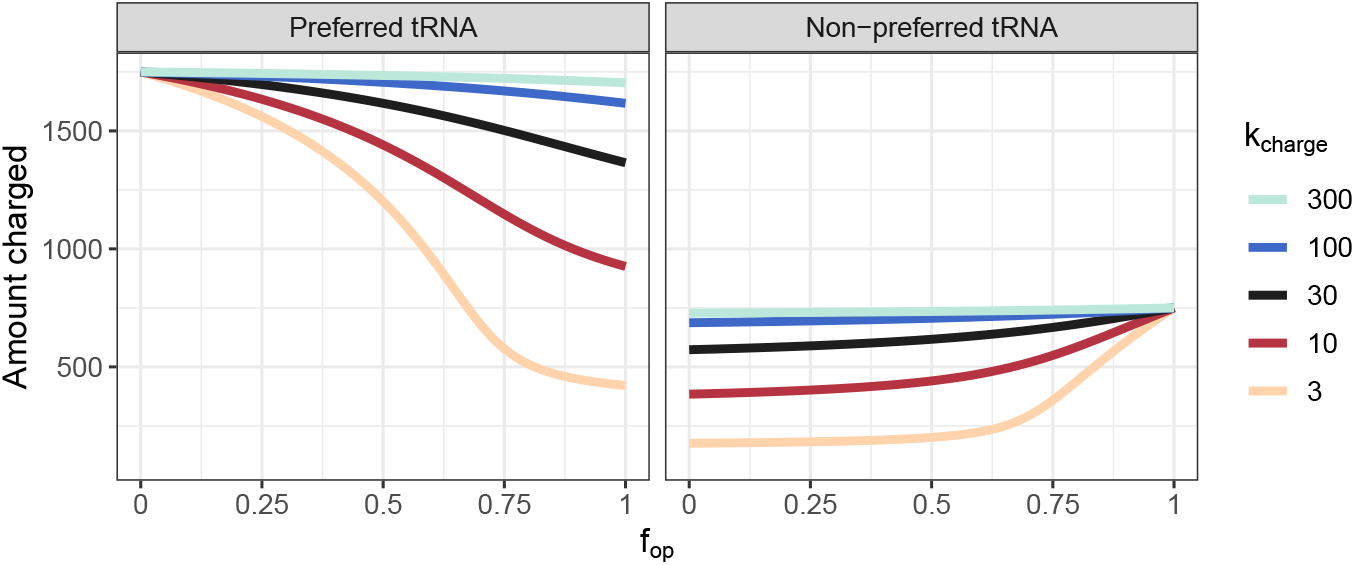
The effects of codon usage on tRNA availability. Steady-state charged tRNA abundances as a function of *f*op in a system where the fraction of preferred tRNAs is 0.7. The tRNA abundances were computed by solving Eqs. (6)–(12) numerically for *T*_c1_ (charged preferred tRNA) and *T*_c2_ (charged non-preferred tRNA).

### A scaling law for total tRNA abundances

Careful inspection of the steady-state equations of the two-codon system [Eqs. (6)–(12)] reveals a useful scaling law for total tRNA abundances. We can arbitrarily increase or decrease the total tRNA amount without changing the resulting protein production rate *P*_r_, by appropriately rescaling the tRNA charging and ribosome elongation constants. This scaling law is helpful for stochastic simulations of the system, as it allows us to run simulations with comparatively low tRNA numbers without having to worry about obtaining unrealistic results.

Assume we rescale the total number of tRNAs in the system by a factor *α*, 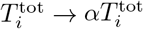. Then, Eqs. (11) and (12) become

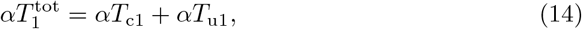

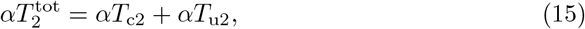

which are equivalent to the original equations.

In Eq. (13), we can compensate for the change in charged tRNA abundances by rescaling the ribosome elongation rate constant *k*_speed_ by 1*/α, k*_speed_ *→ k*_speed_*/α*. Then, Eq. (13) becomes

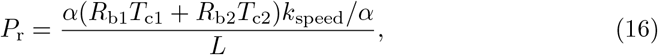

and *α* cancels.

For the remaining three equations involving tRNAs, the *α* terms also cancel. However we now need to account for the re-scaled *k*_speed_ in Eqs. (7) and Eqs. (8). To do so, we rescale *k*_charge_ by 1*/α* as well, so that we have

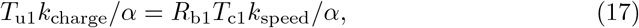

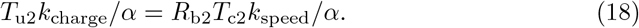

This analysis shows that it is possible to compensate for any arbitrary increase or decrease to the tRNA concentration by dividing the charging and elongation rate constants (*k*_charge_ and *k*_speed_) by the same scaling factor.

To verify the scaling law, we numerically calculated the protein expression rate *P*_r_ using three different values for *α*, applied to a baseline tRNA concentration of 2500 total molecules. We found that indeed *P*_r_ was independent of total tRNA concentration when *k*_charge_ and *k*_speed_ where appropriately rescaled (Figure 4).

**Fig. 4.**
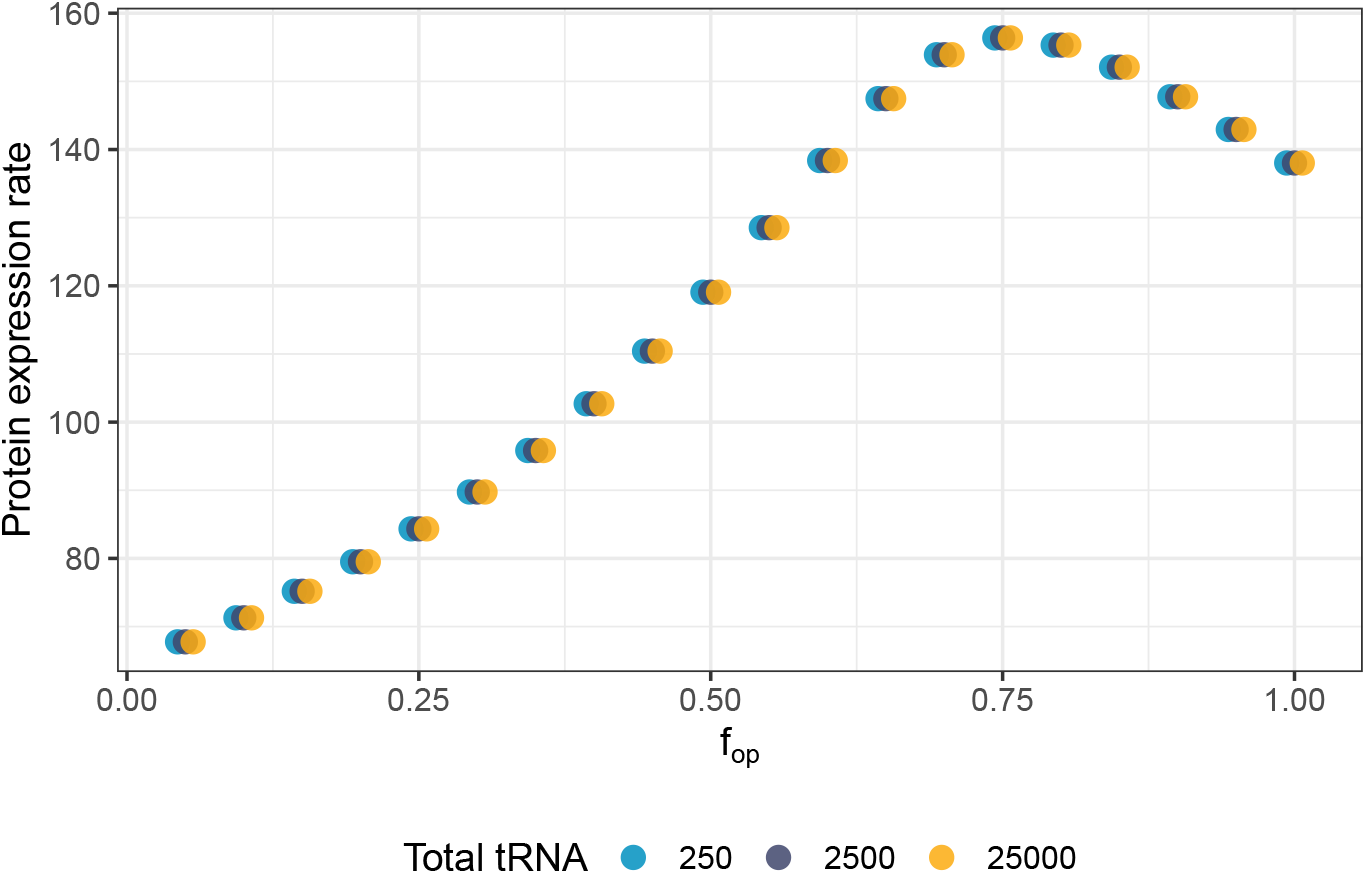
Verification of the scaling law for total tRNA abundances. We used a baseline total tRNA amount of 2500 molecules and *α* values of 0.1, 1, and 10, resulting in tRNA abundances of 250, 2500, and 25,000, respectively. We rescaled *k*_charge_ and *k*_speed_ by *α* to compensate for the change of total tRNA, and then computed the protein expression rate by numerically solving Eq. (13). Since points corresponding to each *f*op overlap entirely, data have been dodged on the horizontal axis to make each overlapping point visible.

### Application to T7 codon deoptimization

Moving towards more biological realism, we used the model with tRNA dynamics to describe gene recoding in the context of bacteriophage T7 gene expression. Bacteriophage T7 has a medium sized (37Kb) genome encoding at least 40 proteins, many of which are responsible for diverting the host bacterium’s cellular resources towards the production of major capsid (10A) proteins (Dunn and Studier 1983; Molineux 2006). Major capsid protein makes up a large portion of the viral envelope that encloses mature T7 particles, making it critical to T7 fitness. The relationship between codon usage in the major capsid gene and T7 fitness has been used to test gene recoding as a viral attenuation strategy for vaccine development. In T7, deoptimization of the major capsid gene caused a nearly 20% fitness reduction (relative to wild-type), measured in viral doublings per hour (Bull et al. 2012). We were interested in whether the large fitness defects might be driven by dynamic depletion of charged tRNAs, caused by suboptimal codon usage in the highly expressed major capsid gene, which had not been systematically explored with a model that incorporates tRNA dynamics.

To simulate codon deoptimization in bacteriophage T7, we combined the tRNA dynamics model with an existing model of T7 gene expression (Jack et al. 2017). Assuming a moderate skew in total tRNA abundances (fraction of preferred tRNAs = 0.7) and low tRNA charging rate (*k*_charge_ = 0.5), we fit the wild-type T7 simulation with tRNA dynamics to experimentally measured T7 gene expression patterns from Jack et al. (2017) (Fig. S2). This was initially done using a small number of tRNA molecules (2500); once the simulation was calibrated, we re-scaled tRNA abundances to a more biologically realistic number (50,000) using the parameter scaling relationship that we had derived earlier (Fig. S3). We were also interested in comparing the bacteriophage simulation with tRNA dynamics to a simulation where the number of charged tRNA molecules is completely static. To make this comparison, we employed an older version of the T7 model where a static translation speed is assigned to each codon to simulate tRNA preference (with lower speeds corresponding to lower cognate tRNA availability). We initially used codon speeds with the same proportions as the tRNAs in the dynamic model, specifically, we used a speed of 7 for the optimal codon, and speed of 3 for the non-optimal codon. Then, we manually adjusted the baseline elongation speed (*k*_speed_) in the static model until both models had similar overall protein expression rates for wild-type gene *10A* codon usage (Fig. S2). To compare the simulations to T7 fitness measurements, we converted simulated capsid protein abundances at 1200 seconds (the time point at which capsid protein makes up the majority of protein in the simulation) to doublings per hour, using a previously reported relationship between major capsid abundance and bacteriophage growth rate (see Methods). Finally, we simulated T7 deoptimization by reducing the fraction of optimal codons (*f*_op_) in gene *10A* in both models, and then plotted predicted/simulated fitness as a function of codon usage (Fig. S4).

We found that deoptimization of gene *10A* caused T7 fitness to decline substantially in both models (Fig. S4), although the decline was sharper for the model with dynamic tRNAs. Since the choice of parameters for the static tRNA model was arbitrarily based on the tRNA proportion for the other model, we reasoned that there might be parameter choices where the static tRNA model more closely matches the experiment. To search the parameter space, we increased the difference between the speeds for optimal and non-optimal codons incrementally until the fitness decline in both models (static and dynamic) was approximately the same (Fig. 5). We found that by imposing a severe codon penalty, where the optimal codon is translated 100 times faster than the non-optmal codon, we could make the fitness decline due to codon deoptimization equivalent in both models.

**Fig. 5.**
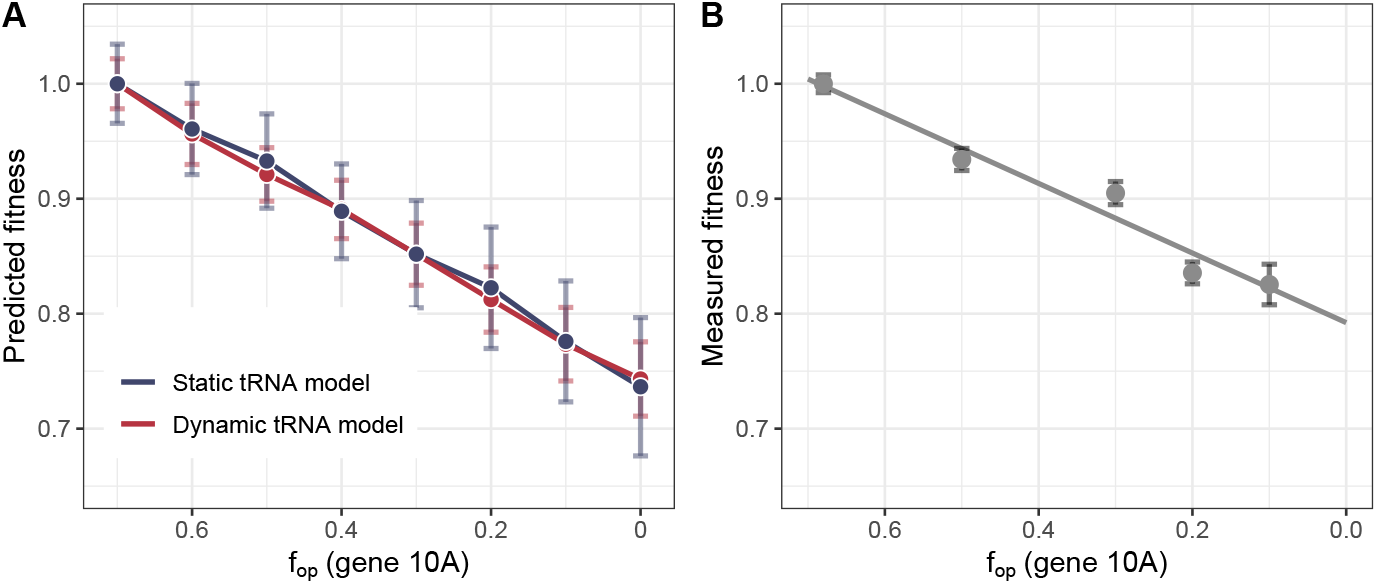
T7 fitness in simulations with tRNA dynamics (dynamic tRNA model) and with static codon speeds (static tRNA model) with a severe codon penalty. A: Simulated T7 fitness as a function of major capsid (gene *10*) codon usage. For each simulation, major capsid (10A) proteins are taken after 1200 seconds of simulation time, and then converted to fitness values (in doubling per hour) using a previously reported relationship (Bull et al. 2011) (see also Methods). Error bars show the standard error from 10 simulation replicates. B: T7 fitness measurements reproduced from Bull et al. (2012). In both A and B, values are normalized to the fitness for wild-type T7 (*f*op *≈* 0.7).

Finally, to further explore differences between the two models, we analyzed how the levels of charged tRNAs in the dynamic model change over the course of T7 infection. We found that at the start of structural gene expression (which includes the major capsid gene), charged fractions of both tRNAs decline rapidly until charged rare tRNAs are completely depleted (Fig. S5). Interestingly, this occurs even in the wild-type (non-recoded) T7 simulation, and appears to be mostly independent of the level codon usage in gene *10A*. Thus, the fitness decline in the dynamic tRNA model is not driven by charged tRNA depletion due to codon usage as we had initially hypothesized. Rather, it is caused by increased demand for the limiting non-preferred tRNAs. Since charged tRNA equilibration occurs rapidly upon the onset of structural gene expression, major capsid protein production is approximately linear with respect to optimal codon usage, and can be replicated with the static tRNA model using a large enough codon penalty.

## 3 Discussion

We have derived a mathematical model for a system with two dynamic tRNA species, based on a prior stochastic simulation of an analogous system introduced in Roots et al. (2025). We have used the numeric solutions of the mathematical model to explore how different amounts of codon usage impact tRNA availability and protein expression. We have found that, in general, the behavior of the model depends on whether the baseline rate of tRNA recharging is fast or slow, relative to translation initiation. Specifically, when tRNA recharging is inefficient, protein expression can be maximized by matching the fraction of optimal codons to the fraction of preferred tRNAs. The more efficient tRNA recharging becomes, the less important are non-optimal codons for maximal translation throughput. We have also used the tRNA dynamics model to augment a whole-cell simulation of bacteriophage T7 gene expression, to model what happens during a T7 infection when the major capsid (gene *10A*) is systematically codon deoptimized. We have found that in the model with tRNA dynamics, charged rare tRNAs are rapidly depleted following the onset of expression of T7 structural genes, and that increased demand for these depleted rare tRNAs results in a decline in T7 fitness.

We used two separate models—one with tRNA dynamics, and another with static codon speeds—to test the hypothesis that deoptimization of the major capsid gene in bacteriophage T7 drives down availability of charged rare tRNAs. We expected that the model with tRNA dynamics would more easily replicate the measured decline in T7 fitness. However, we were ultimately able to fit both models to the experimental data, even though this required assigning a severe translation speed penalty to non-optimal codons in the static tRNA model. Production of proteins that are expressed late in the T7 infection cycle, after charged tRNA fractions reach a steady-state, is largely linearly dependent on codon usage in the dynamic tRNA model and can be approximated using a model where charged tRNAs are completely static. However, this may not hold true for genes expressed earlier during T7 infection, such as the T7 RNA polymerase. Dynamic changes to the tRNA pool could also disproportionately impact endogenous genes with extreme codon usage biases (such as *E. coli* regulatory sequences (Wohlgemuth et al. 2013)), which we did not model in detail in this study. Future work could explore feedback between T7 codon deoptimization and the expression of early T7 genes and/or endogenous *E. coli* genes.

A striking feature in our model is the appearance of a maximum in the protein expression rate that coincides with intermediate levels of optimal codon usage. For this to occur, two separate things must happen simultaneously. First, the tRNA charging rate must be low enough that charged tRNA abundances are sensitive to codon usage. Second, codon usage bias must be high enough that the demand for preferred tRNAs outstrips the rate of tRNA charging. In this regime, the level of charged preferred tRNAs can drop below the level of charged non-preferred tRNAs, such that a role-reversal occurs where preferred tRNAs are actually associated with lower rates of translation.

Our work differs from many existing theoretical models of translation in that we do not assume constant charged tRNA availabilities, which has been a common simplification in the literature. For example, in one approach based on the totally asymmetric simple exclusion process (TASEP) (MacDonald et al. 1968), ribosome movement across a one-dimensional lattice is simulated using fixed translation speeds averaged over groups of codons (Katz et al. 2022; Raveh et al. 2016; Reuveni et al. 2011; Zarai et al. 2016). Another approach is to simulate translation in the context of an entire bacterial or eukaryotic cell, using biologically realistic species abundances. Detailed whole-cell models that take into account codon usage have been developed for *E. coli* (Levin and Tuller 2020) and yeast (Seeger et al. 2023; Shah et al. 2013), but notably, these models only consider individual tRNAs and not tRNA charging.

Although using fixed charged tRNA abundances is common, there are some models that have incorporated tRNA charging dynamics. For example, Brackley et al. (2011) used a TASEP-like model with tRNA dynamics to explore trade-offs between the translation of small numbers of mRNAs with different codon usage biases. By varying the initiation rate for one mRNA species in the model, they observed a similar dynamic pattern where protein expression can increase and then decrease (although, they found that this was driven by ribosome queuing in their model, and not necessarily tRNAs dynamics specifically). In another example, Elf et al. (2003) introduced a mathematical model for the availability of all 40 *E. coli* tRNAs during amino acid starvation. They predicted that charged tRNA fractions should depend both individual tRNA concentrations and codon usage frequencies; we see this also, albeit for a much smaller number of individual tRNAs. Taken together, our study represents a meaningful contribution to the existing body of theoretical work, while demonstrating agreement with existing models that also consider charged tRNA dynamics.

Finally, both the mathematical model and the T7 whole-cell model explored here build on a general purpose simulation framework that has been used to model gene expression dynamics across several different prokaryotic systems (Hill et al. 2024; Jack et al. 2019; Shah et al. 2022). The updated simulation framework with tRNA dynamics is available on github, and could be applied to other systems to make predictions about the impact of codon usage and tRNA availabilities on protein production.

## 4 Methods

In this study, we used a mixture of numerical methods and stochastic simulations to investigate the effects of codon usage bias and tRNA dynamics on prokaryotic gene expression. All simulations were performed using the Pinetree stochastic gene expression simulation software (Jack and Wilke 2019), development version 0.4.1. The code required to reproduce all results is available on GitHub (https://github.com/alexismhill3/tRNAdynamics).

### 4.1 Stochastic simulations of the two-codon model

To verify the numeric solutions to Eqs. (6)–(13), we re-ran the stochastic simulations from Roots et al. (2025) that had been used to simulate cell burden under codon deoptimization. These simulations were set up with the counts for all molecular species (ribosomes, mRNAs, and tRNAs) scaled down by several orders of magnitude relative to a reference bacterial cell. Specifically, the stochastic model used 120 total mRNA molecules, 500 ribosomes, and 2500 tRNAs (for reference, a cell growing under normal laboratory conditions contains 5-10 times as many ribosomes and mRNAs, and 5-10 times as many tRNAs as ribosomes (Bartholomäus et al. 2016; Bremer and Dennis 2008; Dong et al. 1996; Mackie 2013)). These simulations also used two different types of transcripts to represent an exogenous mRNA and an endogenous cellular mRNA, respectively. In our simulations, we included only 100 copies of the endogenous mRNA. All other simulation parameters were exactly as described (Roots et al. 2025).

### 4.2 Numerical solutions of the two-codon model

To solve the two-codon model numerically, we parameterized Eqs. (6)–(12) using the values in Table 2 and then solved for the steady-state number of bound ribosomes (*R*_b1_ and *R*_b2_) and charged tRNAs (*T*_c1_ and *T*_c2_) with different amounts of optimal codon usage (*f*_op_). We used the nsolve() function from the Sympy package in Python (https://www.sympy.org/en/index.html) with the modified newton solver option (solver=“mnewton”) to perform this computation, and then used Eq. (13) to compute the protein expression rate (in molecules per second) from the steady-state numerical solutions. To determine the *f*_op_ that maximizes protein expression 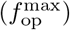, we use the max() function from Pandas to compute (for each set of models, grouped by *k*_charge_ and the fraction of preferred tRNAs) the *f*_op_ corresponding to the maximum point. Since we use discrete values for *f*_op_, the computed maximum may deviate slightly from the true maximum for each curve.

**Table 2.** Summary of parameter choices for the simplified two-codon model.

| Parameter | Value | Units | Description |
| --- | --- | --- | --- |
| $T_{\text{tot}}$ | 2500 | molecules | Total number of tRNAs |
| tRNA ratio | 0.5–0.9 | - | Fraction of preferred tRNAs |
| $R_{\text{tot}}$ | 500 | molecules | Total number of ribosomes |
| $N$ | 100 | molecules | Total number of transcripts |
| $L$ | 300 | codons | Transcript gene length |
| $k_{\text{charge}}$ | 3–300 | $\text{s}^{-1}$ | tRNA charging constant |
| $k_{\text{bind}}$ | $2.5 \times 10^6$ | $\text{M}^{-1}\text{s}^{-1}$ | Ribosome binding constant |
| $k_{\text{speed}}$ | 0.02 | codons $\text{s}^{-1}$ | Ribosome elongation constant |
| $f_{\text{op}}$ | 0.00–1.00 | - | Fraction of optimal codons |

### 4.3 Bacteriophage T7 simulations

The T7 simulations used here expand on a prior whole-cell model of bacteriophage T7 infection dynamics (Jack et al. 2019) that uses the Pinetree simulation software (Jack and Wilke 2019). The model accounts for several major biological events during T7 infections, including injection of the phage genome into the host cell and transition from early to late T7 gene expression, which requires the synthesis of a phage-encoded T7 polymerase and down-regulation of the *E. coli* transcription machinery. The original implementation of the model did not incorporate explicit tRNA dynamics. Instead, tRNA preference was simulated by applying a per-codon ribosome elongation speed multiplier, where lower codon speeds correspond to poor cognate tRNA availability.

We re-ran the T7 simulations using an updated version of the Pinetree simulator that includes tRNA dynamics. First, simulation scripts from the prior T7 modeling project were downloaded from https://github.com/benjaminjack/phage simulation. We then selected the phage_model_recoded.py script which was used to simulate T7 expression with gene *10A* recoding. Since this model had already been calibrated to T7 biology using values derived from prior T7 modeling work and from experiments, we left most preexisting parameters unchanged, adjusting only the baseline ribosome elongation speed (*k*_speed_) as needed to fit the model to T7 gene expression measurements from Jack et al. (2019). The final parameters chosen were a moderate skew in total tRNA abundances (fraction of preferred tRNAs = 0.7), a tRNA charging rate of 0.5, (*k*_charge_ = 0.5), and baseline ribosome elongation speed of 1.5 (*k*_speed_ = 1.5). We also incorporated a function to randomize the placement of optimal and non-optimal codons in the recoded major capsid gene. We ran all simulations for 1200 seconds, corresponding to the point in the T7 infection cycle when gene expression is dominated by synthesis of major capsid proteins and cells begin to lyse. To obtain simulated T7 fitness values, we used *10A* protein abundances after 1200 seconds from 10 independent simulation replicates.

To compare the updated T7 model with tRNA charging to simulations where tRNA availabilities are not dynamic, we re-calibrated the original model with static codon speeds using a similar procedure. Using codon-specific speeds with the same proportions as the tRNA abundances in the dynamic model, we adjusted the baseline elongation speed (*k*_charge_) in the static tRNA model so that the final capsid protein abundances in *both* models (specifically, the wild-type version without any codon deoptimization) were roughly equal to each other. The final parameters used were *k*_speed_ = 2.5 and codon speeds of 7 and 3 for the optimal and non-optimal codon, respectively. We also simulated T7 codon deoptimization using a model with a more severe codon penalty, specifically, a speed of 100 for the optimal codon and 1 for the non-optimal codon. For the model with a severe codon penalty, we needed to increase *k*_speed_ to 5 in order to preserve the wild-type protein abundances across all models.

### 4.4 Converting simulated protein abundances to fitness values

To compare T7 simulations to fitness measurements (reported in viral doublings per hour) we followed the same procedure as was done in previous T7 modeling work (Jack et al. 2019). Briefly, we used a relationship that relates the burst size (*b*) of an infection to the intrinsic growth rate (*r*) (Bull et al. 2011):

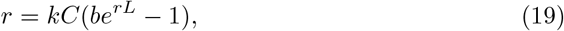

where *k* is the absorption rate, *C* is cell density, and *L* is lysis time. As in prior work, we assumed a lysis time of 12 minutes, a virion size of 400 capsid proteins, and a constant absorption rate and cell density of 1/min, so that Eq. (19) becomes

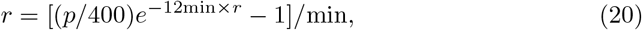

where *p* is the number of major capsid proteins in a simulation after 1200 seconds. We solved for *r* numerically using the uniroot() function in R. Finally, we converted the intrinsic growth rate to doublings per hour *d* using the following equation:

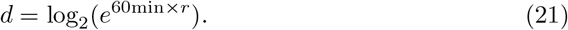

## 5 Data and Code Availability

All analysis and simulation code is available on GitHub (https://github.com/alexismhill3/tRNAdynamics).

## 6 Acknowledgments

The Texas Advanced Computing Center (TACC) at The University of Texas at Austin provided computational resources that have contributed to the research results reported within this paper.

## Supplementary Information

**Supplementary Figure S1.**
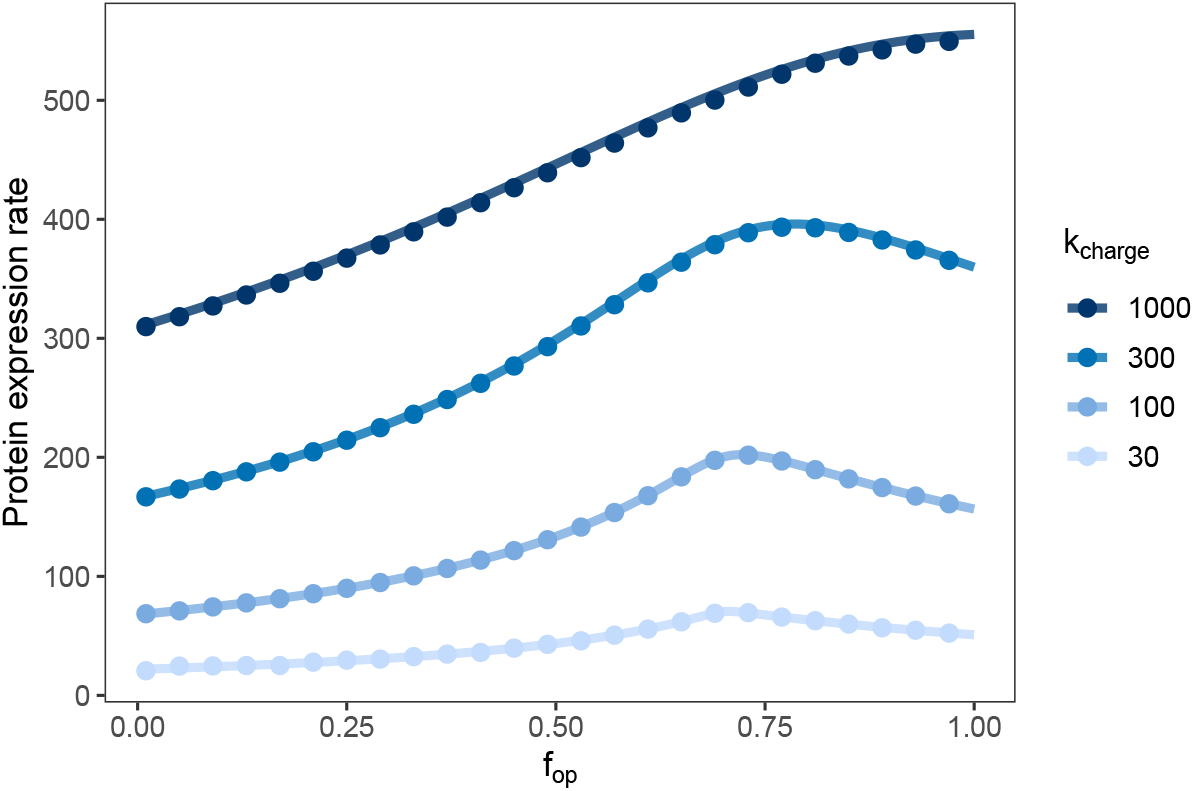
Agreement between simulations and numeric solutions of the two-codon system. In this system, a singular protein product is produced with a rate that depends on the fraction of optimal codons, *f*_op_, and the tRNA charging rate, *k*_charge_. Points show the protein expression rate (in molecules per second) in simulations of this system. Lines show the protein expression rate computed by solving Eq. (13) numerically using parameters that are identical to the simulations. For these simulations, we allow each to reach a steady-state equilibrium and then take the change in total protein abundance over an interval of one second simulated time; points show the average protein expression rate from three simulation replicates.

**Supplementary Figure S2.**
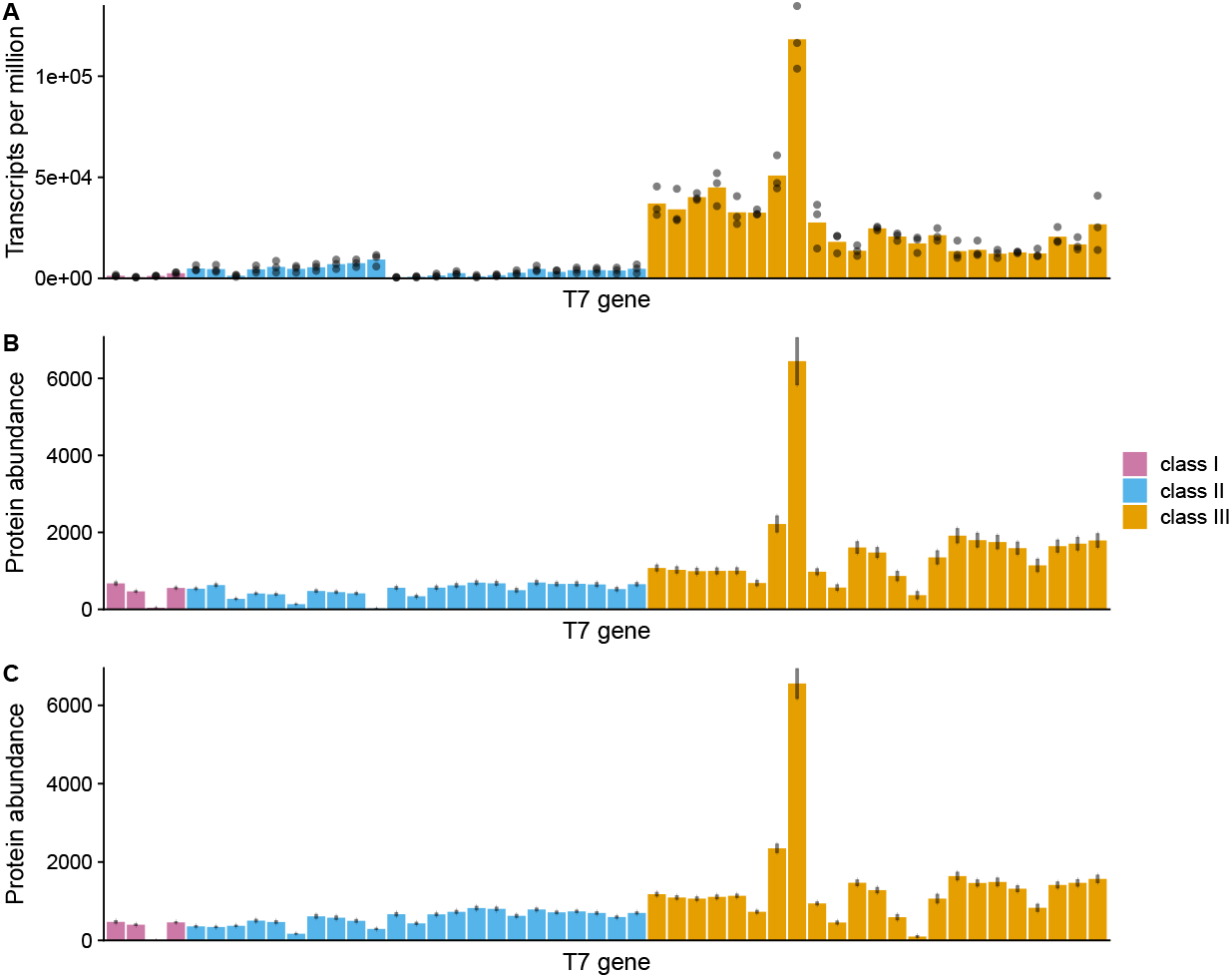
Simulated and measured wild-type T7 gene expression patterns. A: Relative transcript abundances for all T7 genes at 9 min postinfection, just before cell lysis begins, as previously shown in Jack et al. (2019). Each colored bar represents one gene, and genes are arranged from left to right in the order in which they appear in the T7 genome. B and C: Simulated protein abundances in the static tRNA model (B) and dynamic tRNA model (C) after 1200 seconds of simulated time. Error bars show the standard error from 10 simulation replicates.

**Supplementary Figure S3.**
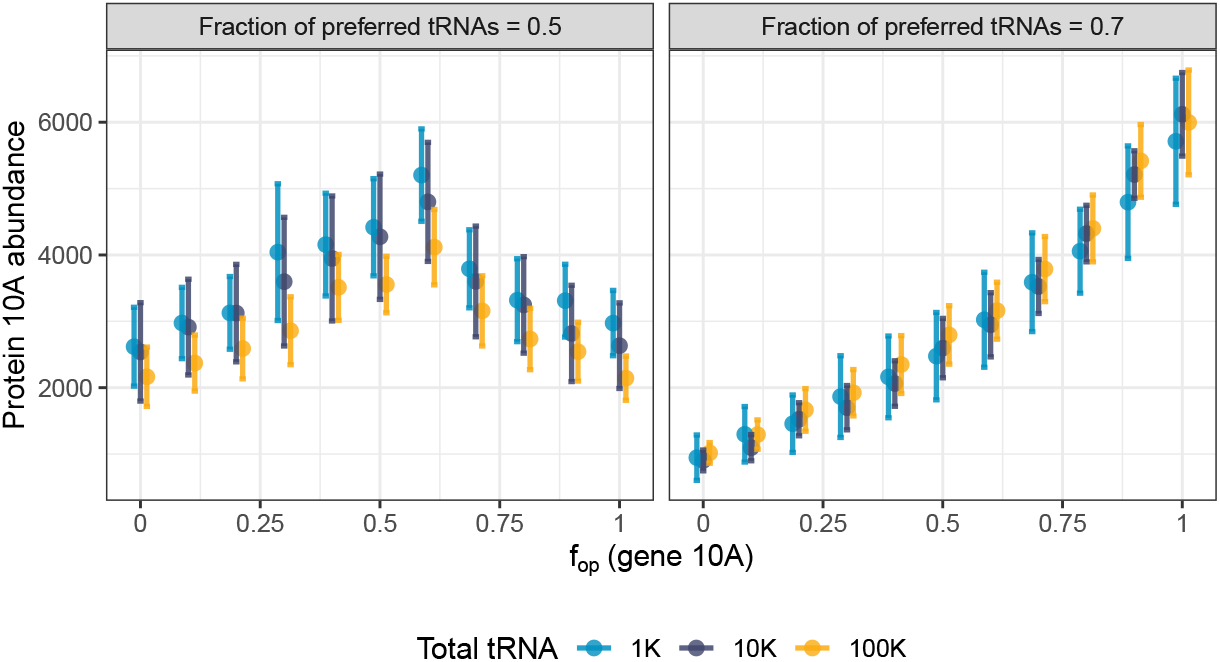
Scaling law for tRNA abundances remains valid in T7 simulations. We ran three sets of T7 simulations with the total number of tRNAs scaled by a factor of 0.1, 1, and 10 relative to the baseline tRNA concentration of 10,000 molecules, while correspondingly rescaling *k*_charge_ and *k*_speed_. Shown are the resulting protein 10A/major capsid protein abundances after 750 seconds of simulation time as a function of the fraction of optimal codons (*f*op) in gene *10A*. The error bars indicate the standard error from three independent simulation replicates. We observe that the tRNA scaling law remains approximately correct in the more complex T7 simulations.

**Supplementary Figure S4.**
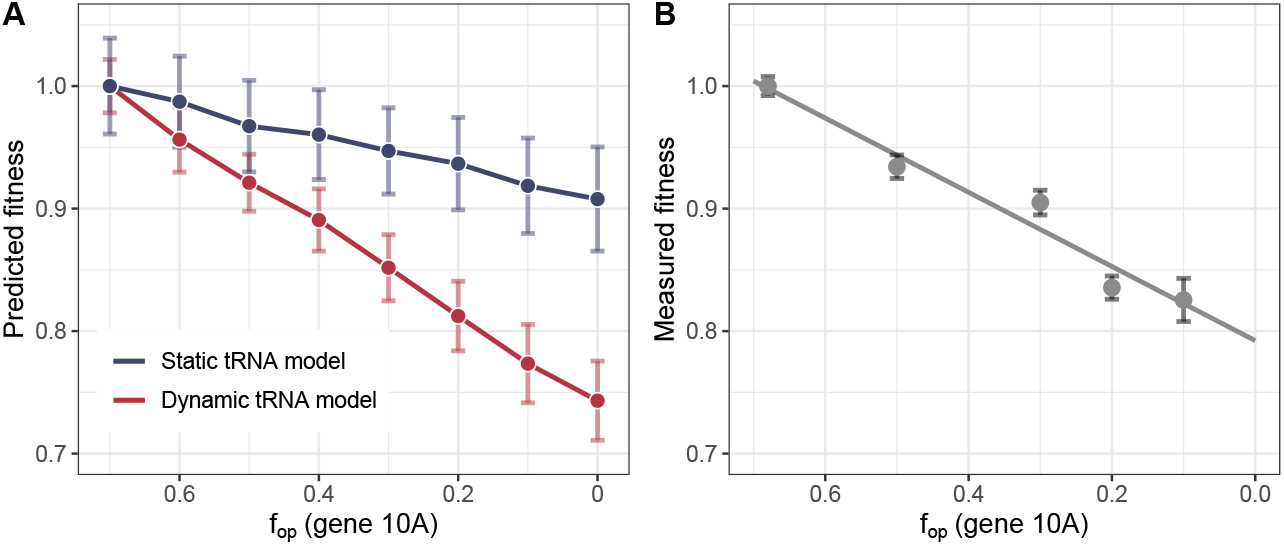
T7 fitness in simulations with tRNA dynamics (dynamic tRNA model) and with static codon speeds (static tRNA model) that match tRNA proportions in the dynamic tRNA model. A: For the dynamic tRNA model, we used a moderate skew in total tRNA abundances (fraction of preferred tRNA = 0.7). For the static tRNA model, we used codon speeds of 7 and 3 for preferred and non-preferred codons, respectively. Error bars show the standard error from 10 simulation replicates for each model. B: T7 fitness measurements reproduced from Bull et al. (2012). In both A and B, values are normalized to the fitness for wild-type T7 (*f*op *≈* 0.7)

**Supplementary Figure S5.**
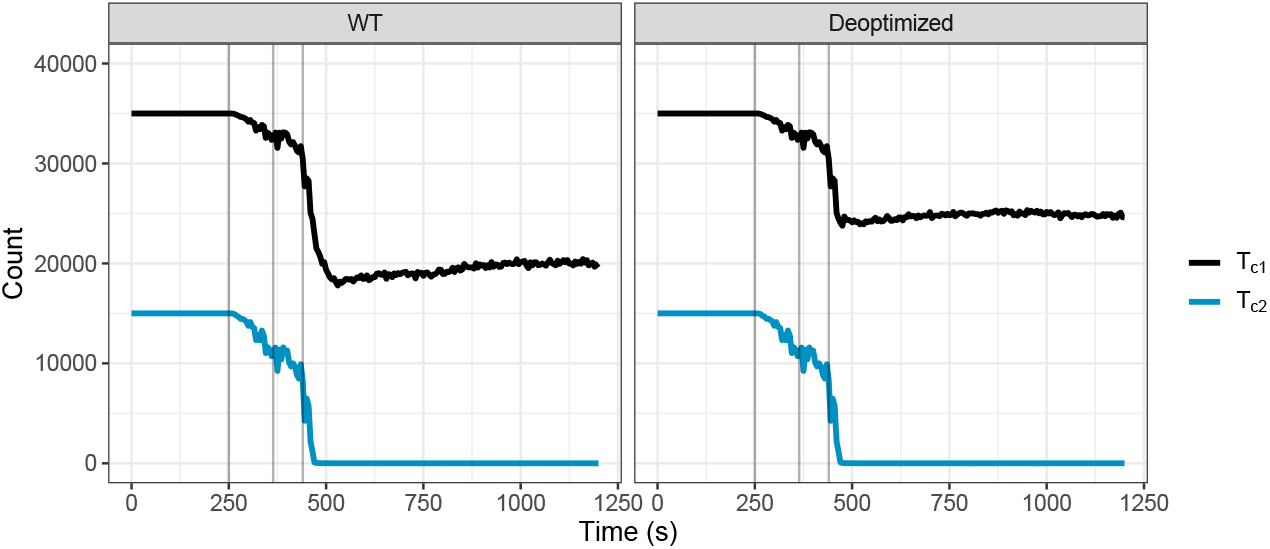
Charged tRNA abundances in simulations of wild-type (WT) and codon deoptimized T7 (gene *10A f*op = 0.1). Total charged tRNA abundances are plotted versus simulation time for *T*_c1_ (charged preferred tRNA) and *T*_c2_ (charged non-preferred tRNA). The gray vertical lines indicate notable simulation events with respect to T7 biology. From left to right (in each panel): first appearance of T7 transcripts, first appearance of T7 RNA polymerases, and beginning of transcription of structural genes.

